# Microtubule growth rates are sensitive to global and local changes in microtubule plus-end density

**DOI:** 10.1101/849190

**Authors:** Zachary M. Geisterfer, Daniel Zhu, Timothy Mitchison, John Oakey, Jesse C. Gatlin

## Abstract

The microtubule (MT) cytoskeleton plays critically important roles in numerous cellular functions in eukaryotes, and it does so across a functionally diverse and morphologically disparate range of cell types [1]. In these roles, MT assemblies must adopt distinct morphologies and physical dimensions to perform specific functions [2-5]. As such, these macromolecular assemblies—as well as the dynamics of the individual MT polymers from which they are made—must scale and change in accordance with cell size and geometry. As first shown by Inoue using polarization microscopy, microtubules assemble to a steady state in mass, leaving enough of their subunits soluble to allow rapid growth and turnover. This suggests some negative feedback that limits the extent of assembly, for example decrease in growth rate, or increase in catastrophe rate, as the soluble subunit pool decreases. Such feedbacks might be global or local. Although these ideas have informed the field for decades, they have not been observed experimentally. Here we describe an experimental system designed to examine these long-standing ideas and determine a role for MT plus-end density in regulating MT growth rates.

## Main

MT based assemblies, specifically interphase MT asters and mitotic spindles, have been confined to and characterized in discrete droplets of cell-free Xenopus egg extract [6-9] to study MT self-organization and scaling phenomena. In these droplets, however, it is difficult to collect images with the sufficiently high spatial and temporal resolution needed to characterize dynamic molecular-scale phenomena such as MT growth. This is partly due to the propensity of droplets to fuse, and the unpredictable movements of the droplets within the imaging plane. In addition, the spherical geometry of the droplets presents a unique imaging challenge, as light emitted from the specimen must travel through multiple refractive indices before being collected by the objective. Additionally, the spherical shape of the droplets means that cellular events of interest are not always confined to a region near the coverslip. To circumvent these limitations in a way that still allows for precise control of extract volume, we used well-characterized hydrogel photolithography methods [10] and developed an approach that enabled us to confine cell-free extracts and MT asters within biologically inert hydrogel enclosures of precise geometrical shape and size (**Figure 1a**).

**Figure 1.**
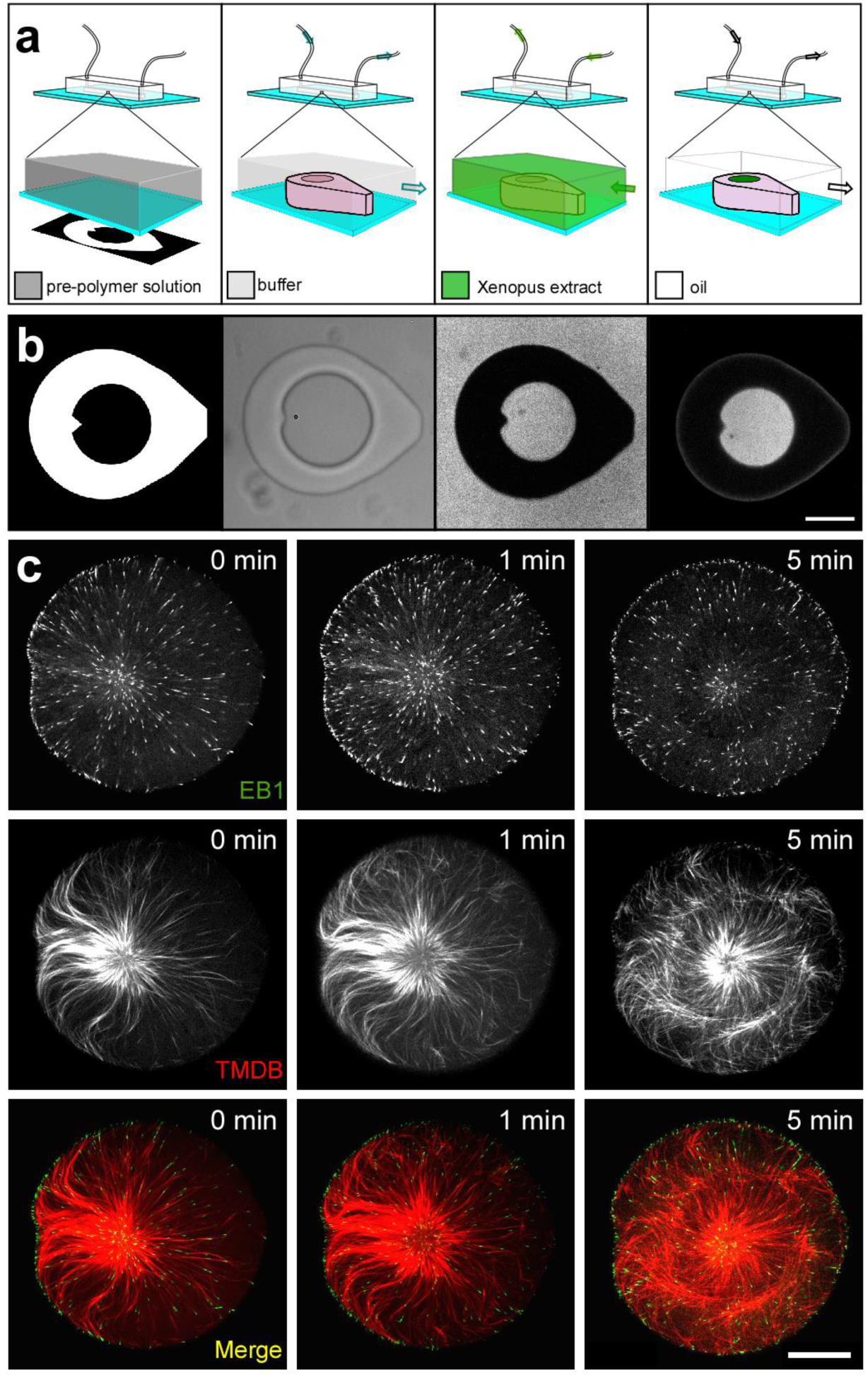
Extract-laden hydrogel micro-enclosures. Schematic **(a)** and captured images **(b)** corresponding to the process for isolating discrete volumes of cytoplasmic extract in hydrogel micro-enclosures. The calibrated digital mask used to create hydrogel structures are pictured in the leftmost panels. Schematic and bright-field image of the micro-enclosure (center-left panels), which is filled with Xenopus extract containing a soluble fluorophore (center-right panels) and then isolated by oil crossflow (rightmost panels) and imaged using fluorescence microscopy (scale bar in b = 50 μm). MT aster formation **(c)** in micro-well enclosures visualized through EB1-GFP and mCherry-TBMD (scale bar = 25 μm). 0 min corresponds to the start of the image series.

To photo-pattern structures on the coverslip surface, we placed a digital micro-mirror array in the light path of a microscope and projected light patterns onto a pre-polymer solution of poly(ethylene glycol) diacrylate (PEGDA) contained within a microfluidic flow chamber (**Figure 1a, b; leftmost and center-left panels**). The positions of these enclosures within the device were dictated by the random spatial arrangement of artificial microtubule organizing centers (aMTOCs) present within the pre-polymer solution. By first generating a flow of extract into the device from one direction and then subsequently generating the flow of an oil phase (fluorinated oil/surfactant) from the opposite direction, we could trap aqueous volumes of extract within the confines of the hydrogel micro-container (**Figure 1a, b**). It should be noted that the exterior tear-drop shape of the hydrogel micro-enclosure was important for isolating extract or other aqueous phases in hydrogel structures using oil crossflow (**Supplementary Figure 1**). With this experimental paradigm, we successfully isolated discrete volumes of cell-free extracts at the coverslip surface (**Figure 1a, b; rightmost panel**) with precise control of geometry and volume. aMTOCs confined in this manner were able to nucleate MTs and generate MT asters as described previously in bulk interphase egg extracts [11, 12] (**Figure 1c**). MT plus-ends and the MT lattice were visualized using EB1-GFP and a fluorescent MT-associated protein, mCherry-TBMD [13], respectively (**Figure 1c**).

Given that the size of macromolecular MT-based assemblies has been shown to scale with changes in the size of the cell in which they are made [6, 7, 14, 15], we predicted that changes in cytoplasmic volume might affect MT growth rates. To test this hypothesis, we used our hydrogel experimental platform to capture aMTOCs in cylindrical micro-enclosures of increasing diameter. The range of diameters tested (equivalent to spherical cells ranging from 30 to 115 µm in diameter [Ø]) was chosen based on the range of blastomere sizes in which *in vivo* mitotic spindle scaling has been observed ([7]; **Figure 2a**). The extracts used in these experiments were supplemented with a low concentration of EB1-GFP (60 nM) to visualize and track growing MT plus-ends. Subsequent analysis of time-lapse recordings of EB1 comets allowed us to plot MT growth rates versus micro-enclosure volume (**Figure 2b; Supplementary Figure 2a**).

**Figure 2.**
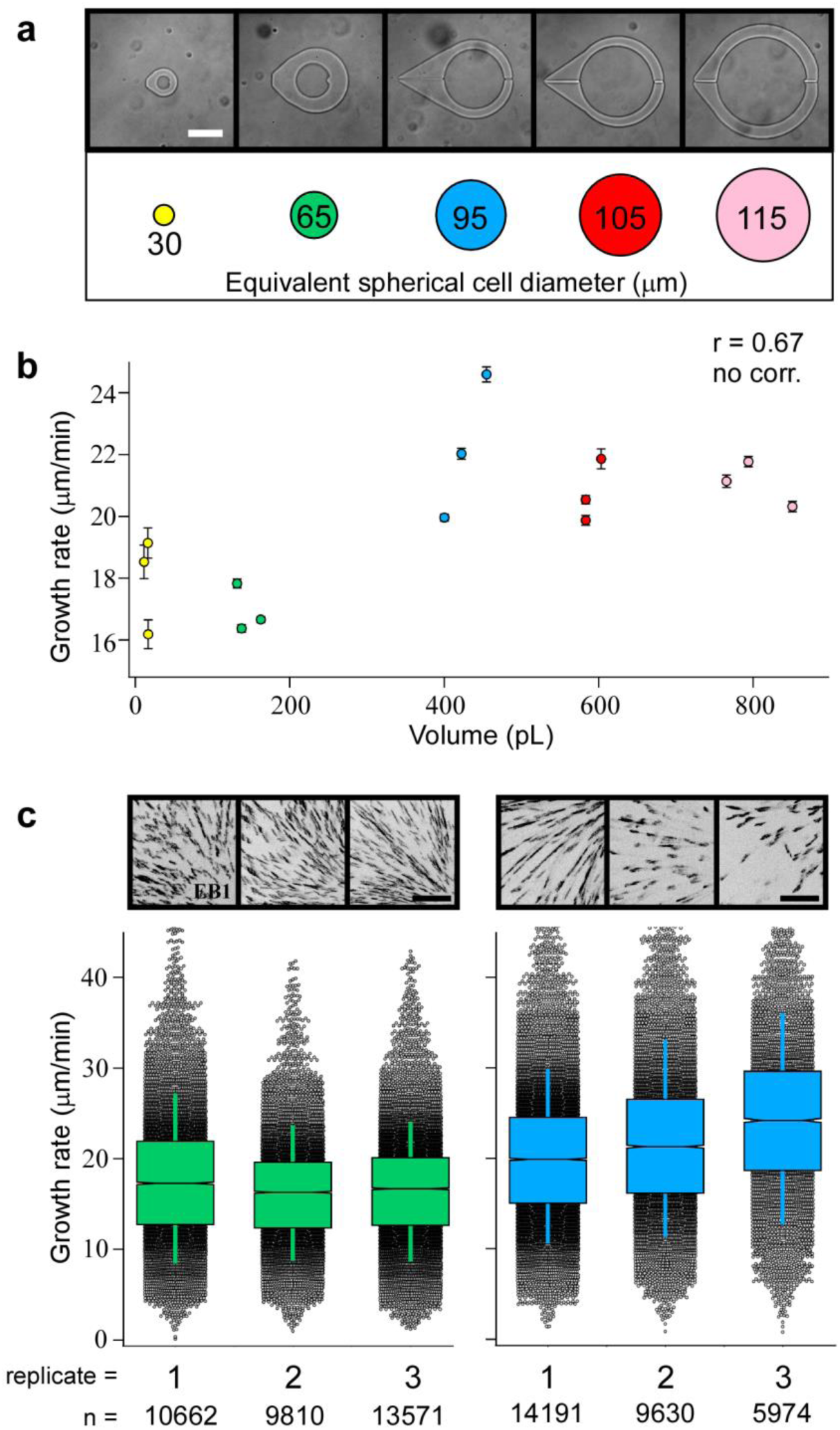
Effects of cytoplasmic volume on MT growth rates. **(a)** Cylindrical micro-enclosures of increasing internal diameters are shown along with their equivalent spherical cell diameters (scale bar = 75 μm). **(b)** MT growth rates plotted as a function of cytoplasmic volume. Error bars equal two SEMs. The Pearson correlation coefficient (r) is indicated at the top of the graph and is significant if p < 0.01 (the absence of a correlation is indicated otherwise). A comparison of MT growth rates **(c)** in the experimental replicates for ∼160-pL (left) and ∼400-pL (right) volumes (∼65 μm and ∼95 μm Ø, respectfully; scale bar = 5 μm). Box plots feature a Tukey-style interquartile range, with whiskers indicating one SD of the median. Notches approximate the 95% confidence interval for the median. Representative max-intensity projections for each imaging series displaying EB1 signal in black are shown above the graphical data. The total number of EB1 tracks (n) recorded for each imaging series are displayed at the bottom for reference.

MT growth rates were, in general, proportional to cytoplasmic volume within our system. However, some of the experimental data were inconsistent with this trend. In volumes as low as ∼13 pL, we observed growth rates consistent with those found in ∼160-pL enclosures (corresponding to spherical cells of ∼30 and 65 µm Ø, respectively), and MT growth rates seemed to plateau at cytoplasmic volumes above ∼400 pL (spherical cells of ∼95 µm Ø; **Figure 2b**). Inexplicably, we also observed large, statistically significant differences in MT growth rates in micro-enclosures of similar volumes (e.g., see the spread observed in the data points for enclosures around 400 pL in volume in **Figure 2b**). These observations suggested that there might be some other unaccounted for variable influencing MT growth rates in this system. Indeed, when we compared EB1-GFP signal densities within each ∼400-pL device, we found large differences in the total number of tracked MT growing ends (**Figure 2c**; **Supplementary movie 1**). In contrast, MT growth rates observed in devices with volumes of ∼160 pL (cell diameter of ∼65 µm) were far more homogeneous, with less overall variation in MT growth rates and EB1 signal density (**Figure 2c**; **Supplementary movie 2**). From these data, we speculated that the differences in MT growth rates observed in similar cytoplasmic volumes might be caused by differences in MT plus-end density, rather than changes in cytoplasmic volume. This inference would be consistent with a scenario in which MT plus-ends act as a sink for both structural components (i.e., tubulin) and/or +TIP proteins [16-19].

To determine whether MT plus-end density is a predictor of MT growth rates within our system, we re-plotted MT growth rates versus EB1 density (**Figure 3a; Supplementary Figure 2b**). Over the range of EB1 comet densities, MT plus-end density negatively correlated with MT growth rates and could account for the device-to-device variation found in MT growth rates in near-isovolumetric enclosures (**Figure 3a; Supplementary Figure 2b**). These data suggest that growth rates are perhaps regulated in a manner in which growing ends compete with one another for a limited supply of components or local activation by key factor(s) [20, 21]. By this logic, we hypothesized that changes in MT plus-end density would lead to correlative changes in MT growth rates. To test this, we increased MT plus-end density by capturing two aMTOCs in ∼160-pL micro-enclosures (**Figure 3b; Supplementary movie 3**). With the additional aMTOC, we indeed observed greater densities of EB1 comets, which resulted in measured decreases in MT growth rates (**Figure 3a**). This observation confirmed that MT growth rates are sensitive to changes in global MT plus-end density. We reasoned that the link between MT plus-end density and MT growth rates might depend on the availability of key microtubule associated proteins (MAPs) and their diffusion toward MT growing ends [17, 22].

**Figure 3.**
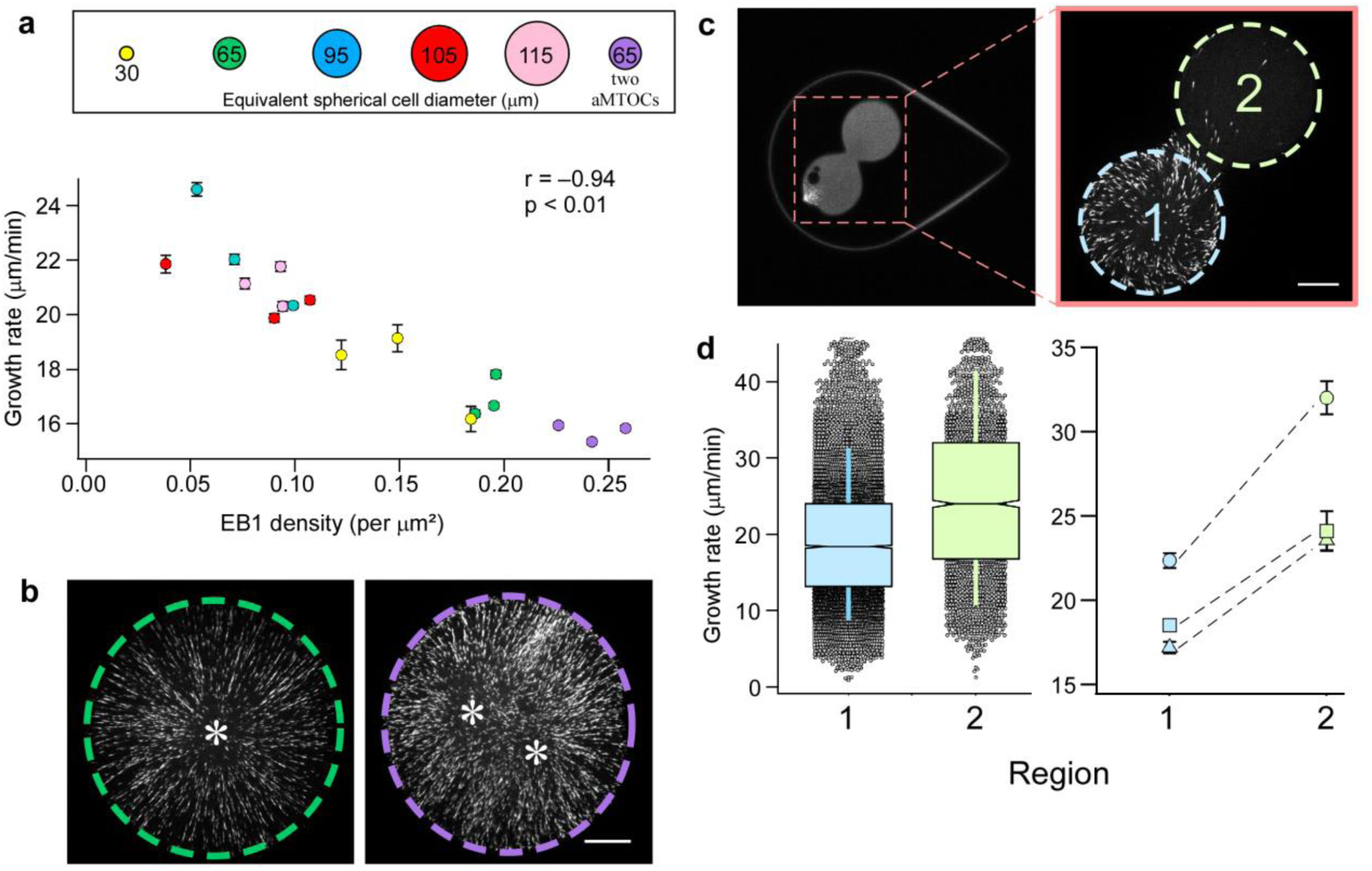
MT growth rates as a function of MT plus-end density. **(a)** MT growth rates displayed as a function of EB1 comet density. Error bars equal two SEMs. Pearson’s correlation coefficient (r) is indicated at the top of the graph and is significant if p < 0.01. **(b)** The effect of an additional aMTOC, with representative images showing EB1 signal from captured time-lapse series. The left micro-enclosure (green dashed line) has one aMTOC, whereas the right micro-enclosure (purple dashed line) has two. Quantified MT growth rates are displayed in the graph in (a). Asterisks denote the relative positions of the aMTOCs (scale bar = 15 μm). **(c)** Hourglass-shaped micro-enclosure. The area indicated in the left panel is shown at higher magnification in the right panel and includes the two lobes of the hourglass enclosure (region 1 in light blue and region 2 in light green; scale bar = 15 μm). The left and right images were acquired from two different hourglass-shaped micro-enclosures. **(d)** MT growth rates from the two regions of the hourglass micro-enclosures. The left graph displays grouped MT growth rates from the two regions of the micro-enclosure as box plots featuring a Tukey-style interquartile range (IQR), with whiskers indicating one SD of the median. Notches approximate the 95% confidence interval of the median for three different micro-enclosures. The right graph shows paired MT growth rates from the two regions of the hourglass micro-enclosure for three different enclosures. Error bars equal two SEMs.

If MT growth rates are diffusion limited in our system, one would expect spatial differences in MT plus-end densities to impart a local effect on MT growth rates. To test this, we generated hourglass-shaped micro-enclosures and trapped an aMTOC in one of the two connected lobes (**Figure 3c**). In each of these experiments, the lobe containing the aMTOC exhibited a much higher EB1 comet density than the unoccupied lobe. A comparison of MT growth rates in each lobe showed significant differences in MT growth rates (**Figure 3d**), despite all growing ends being contained within the same continuous cytoplasm. Consistent with our previous findings, the regions containing lower MT plus-end densities displayed the highest MT growth rates (**Figure 3d**). This result suggested a possible role for a diffusion-limited mechanism in regulating MT growth rates [17, 23].

The observed variation in MT growth rates within a single cytoplasm led us to explore the length scales over which MT growth rates might be sensitive to spatial differences in MT plus-end density. To identify the local plus-end density experienced by each EB1 comet over its lifetime, we developed a custom MATLAB code that uses positional data obtained from u-track software [24, 25]. This code allowed us to define a search area projected from the center of each comet and then to identify and count the number of neighboring EB1 comets found within that area over the lifetime of the tracked comet (**Figure 4a**). This analysis, termed “local plus-end density”, was then repeated for all tracked EB1 comets in the image series.

**Figure 4.**
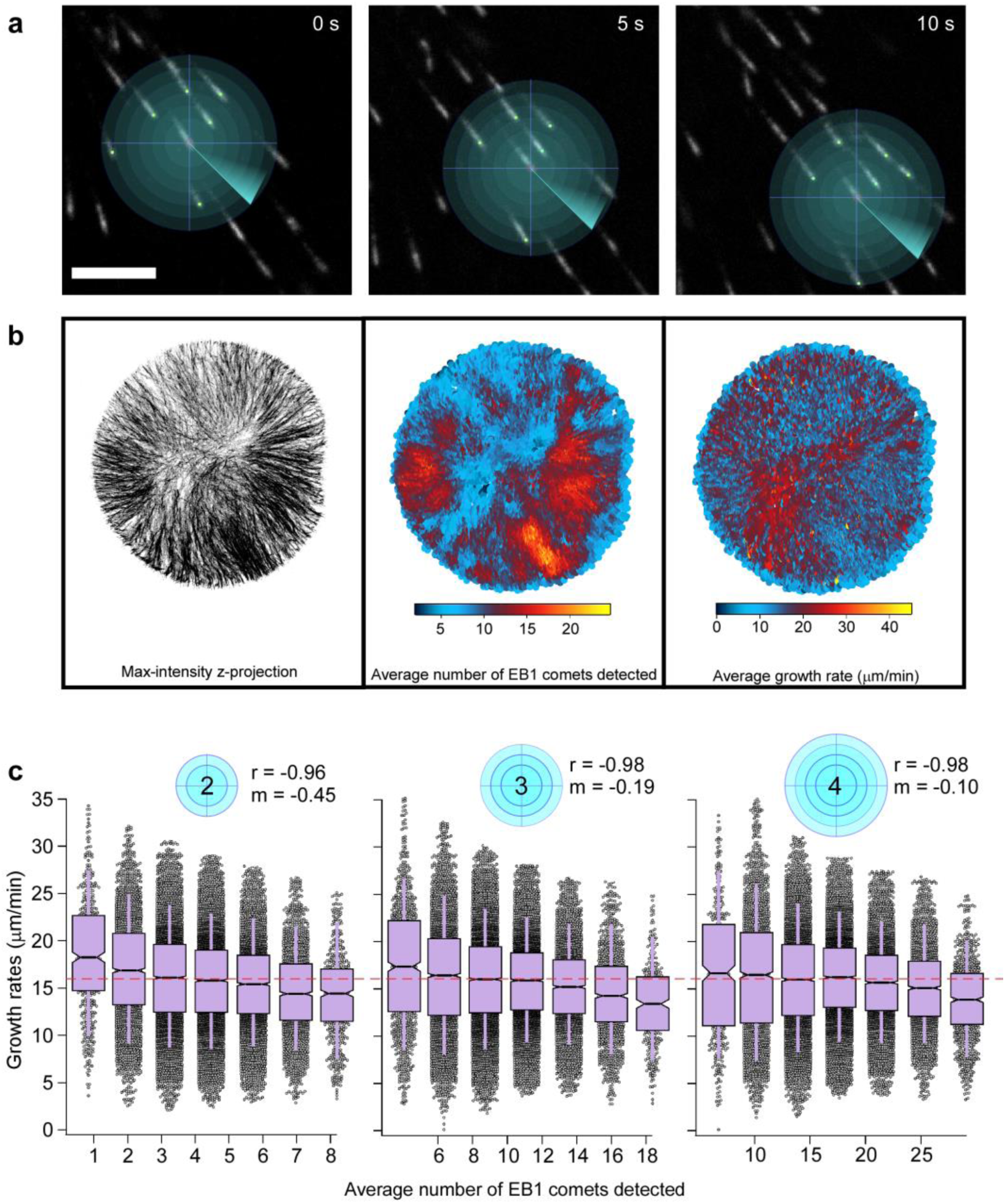
MT growth rates as a function of local MT plus-end density. Pictorial representation **(a)** of local density analysis, showing a series of search radii projected from the center of an EB1 comet over its lifetime (scale bar = 5 μm). **(b)** A micro-enclosure supplemented with two aMTOCs depicted as a max-intensity z-projection (left panel). The average velocity and average density of each EB1 comet plotted at each coordinate position of the comet’s lifetime (center and right panel, respectively). **(c)** Average density as a function of the search radius (indicated by blue circles), displayed as a box plot with bin sizes being equal to a 10% increment of the maximum density observed at that search radius (red dashed line indicates global density of the micro-enclosure). Box plots feature a Tukey-style IQR, with whiskers indicating one SD of the median. Notches approximate the 95% confidence interval of the median.

We first analyzed the relationship between local plus-end density and MT growth rates in a time-lapse image series in which spatial variation in growing plus-end density was qualitatively evident (one of the “two aMTOCs” time series in **Figure 3a, b**; see also **Supplementary movie 3**). This allowed us to compare regions of varying plus-end density at similar radial distances from the aMTOC. Moreover, the simple cylindrical geometry of the extract as confined in this device allowed us to rule out the possibility that spatial signaling gradients emanating from the aMTOC might contribute to the differences in MT growth rates observed in our two-lobed devices (**Figure 3c, d**). After determining the local plus-end density experienced by each EB1 comet, we plotted the average local density and average growth rate for each EB1 comet over its lifetime against the x-y position (center and right panel, respectively, **Figure 4b**). Consistent with the negative correlation we observed between global plus-end density and mean MT growth rate (**Figure 2a**), regions with lower MT growth rates corresponded to those regions showing higher local MT plus-end densities (**Figure 4b;** see also **Supplementary figure 3**). This negative relationship could be seen at multiple positions and distances from the nucleation source, suggesting that the local MT plus-end density was indeed a predictor of individual MT growth rates.

To better characterize this relationship and determine the length scale over which the local density could influence individual MT growth rates, we performed the local density analysis over a range of search radii from 2 to 4 μm. The output from each search radius was binned and plotted, with each bin centered on the average local density contained within that bin (see **Supplementary Figure 4**). This analysis revealed that MT growth rates were negatively correlated with local MT plus-end density (**Figure 4c**). Not surprisingly, at smaller search radii, the absolute value of the slope of the linear fit increased, suggesting that the effect of MT plus-end density on MT growth rates is more pronounced at smaller distances (**Figure 4c**). Moreover, as the search radius increased, the slope of the linear fit approached zero, and the average velocity of each bin reflected the mean global growth rate of the micro-enclosure (red dashed line, **Figure 4c**). This relationship between local MT plus-end density and MT growth rates was observed across a range of cytoplasmic volumes (**Supplementary Figure 5**), suggesting that diffusion of components, rather than the absolute protein content of the cytoplasm, is responsible for the observed local density effect.

Our analyses indicated that MT growth rates in confined cytoplasm decrease concomitantly with increases in MT plus-end density (**Figures 3, 4**). Mechanistically, these results suggest (i) that each MT growing end might act as a local sink for either tubulin or key regulators of MT growth (MT plus-end competition), or (ii) that steric hindrance and changes in viscosity within dense MT polymer networks impede the loading of components onto MT plus-ends [23, 26]. Mechanism (i) requires either a limited source of structural components or the limited translation or rotational diffusion of these same key elements. Predictions for mechanism (ii) are less clear, with crowding effects that are due to large crowding agents (BSA, PEG, etc.) speeding up chemical reactions through an “excluded volume effect”, whereas small crowding agents (ethylene glycol, glycerol) slow down the same reactions in a diffusion-limited manner [23, 26, 27].

In summary, our analyses suggest that a steady-state dynamic MT assembly can experience a global depletion of components, whereas an individual growing MT end experiences a local component gradient dictated by the presence and proximity of other growing MT ends. We speculate that these density-dependent effects likely exist because of constraints imposed by the slow diffusion of MT growth–promoting factors. The case could be made for tubulin as one such factor; however, Odde predicted that tubulin concentration would return to its bulk concentration at only ∼50 nm from a growing MT end [16], which is well below the length scales of the density-dependent effect we have identified. Though the *in extract* conditions used here differ considerably from Odde’s conditions, we still argue that tubulin is most likely not limiting MT growth rates in our system based on the protein’s relatively large diffusion coefficient and high concentration (estimated to be ∼15–20 μM in extracts [28]). Rather, the slower rotational and translational diffusion of other larger, and less abundant, MT growth regulators, e.g., XMAP215 (120 nM [22, 29, 30]), suggests that better candidates exist. In evoking a local sink/limiting component model to explain our observations [31], we have implicitly assumed that changes in other MT dynamic parameters, more specifically those that might contribute to an increase in available tubulin in a compensatory way, are unaffected by MT plus-end density. This assumption will ultimately have to be validated.

This new insight into the regulation of MT dynamics can explain, in part, the high variability in MT growth rates as measured within the same organism [12, 32, 33]. Measurements of MT growth rates both *in vivo* and *in vitro* also vary markedly, with the variability in growth rates far exceeding the ∼2% variability that would be expected from a simple Poisson model of tubulin addition to the MT plus-end ([34]; we measured as high as 21%; unpublished observation). Explanations for this variation have been numerous and experimentally intractable, as many reasonable, and confounding, causes have been evoked, e.g., proximity to membranes, spatial effects resulting from changes in MAP function and post-translational modifications, and steric crowding effects [23, 35, 36]. Our results suggest that some of this variability can be accounted for simply by spatial differences in growing MT plus-end density. Our results show specific subpopulations of MTs as having growth rates that vary from the mean growth rate of the system by as much as ∼17% (**Figure 4c**). In some cases, these marked differences in MT growth rates are brought about by the presence or absence of as few as about seven MT growing ends within a ∼13-μm^2^ area (**Figure 4c**; 2-μm search radius).

In summary, our observations suggest that MT growth rates are spatially regulated by the presence and proximity of other MT plus-ends, and that this spatial regulation can impart local changes in the dynamics of MT subpopulations within a single, continuous cytoplasm. This might explain recently observed differences in astral MT and spindle MT growth rates in *Caenorhabditis elegans* embryos [14]. We postulate that this mechanism might be biologically significant in several contexts, most intriguingly in the context of mitotic spindle size scaling during development, as relatively small changes in MT growth rates have been shown to correlate with large changes in mitotic spindle size [22, 29].

## Methods

### CSF-arrested Xenopus egg extract preparation

Crude Xenopus egg extracts were prepared as described [37]. These CSF-arrested extracts were then induced into interphase 1 h prior to imaging via addition of Ca^2+^ to a final concentration of 400 μM. Translation was inhibited in these extracts 1 h prior to imaging through the addition of cycloheximide to a final concentration of 355 μM. For all conditions where MT growth rates were quantified, extract was supplemented with 60 nM EB1-GFP.

### Preparation of microfluidic flow chambers

Preparation of microfluidic flow chambers was carried out as described [38], with the following modifications. The coverslips used for the preparation of the chambers were first soaked in 1M HCl at 50°C for 16 h before being extensively rinsed with double distilled water. After acid washing, the coverslips were then rinsed with 100% ethanol and left to dry between sheets of Whatman filter paper. In addition, the single-use microfluidic devices cast in PDMS feature a flow channel of ∼3 mm in width and ∼45 mm in length, with a height of 30 μm, as opposed to a T-junction.

Microfluidic flow chambers were PEGylated to ensure a biologically inert glass surface. This was accomplished by first treating the microfluidic flow chamber with 2% (v/v) 3-(trimethoxysilyl)propyl methacrylate in 95% ethanol for 15 min after plasma treatment. The chambers were then flushed with 30 channel volumes of 95% ethanol and heated at 70°C for 15 min. After an additional flush with ddH_2_O, these devices were left to dry at room temperature. These devices were then filled with a solution containing 10% (w/w) poly(ethylene glycol) methacrylate and 0.1% (w/w) lithium phenyl-2,4,6-trimethylbenzoylphosphinate. Each device was then exposed for 3 seconds to 10 mW of UV light (352 nm) at a distance of 15 cm on a reflective surface. After exposure, the device was then immediately flushed with five channel volumes of the same solution and exposed once more. This process was repeated for three exposures in total. After PEGylation, the flow chambers were flushed with 30 channel volumes of ddH_2_O and left to soak in ddH20 at 4°C until use.

### Microfluidic encapsulation of Xenopus egg extract

To create PEGDA micro-enclosures, passivated microfluidic flow chambers were filled with a solution containing 20% (w/w) PEGDA 700, 0.5% (w/w) lithium phenyl-2,4,6-trimethylbenzoylphosphinate and aMTOCs in CSF-XB buffer. The devices were then placed on an IX81 stand (Olympus) equipped with a digital micromirror device (DMD; Polygon 400dense from Mightex). With the use of calibrated digital masks projecting micro-enclosure negatives, the solution within the device was then selectively exposed to 405-nm light until full gelation of the micro-enclosure wall was observed. It should be noted that exposure conditions varied slightly per device because of inconsistencies with the coverslip orientation with respect to the light path. Unused polymer solution was then flushed out of the channel using 30 channel volumes of CSF-XB buffer pumped in through Tygon microbore tubing (0.010-inch ID × 0.030-inch OD; Saint-Gobain Performance Plastics) at a continuous rate of 7 μl/min. Fluid flow to the device was established using a syringe pump (neMESYS, CETONI GmbH). The channels containing PEGDA micro-enclosures were then stored in CSF-XB buffer for at least 10 h prior to filling the device with Xenopus extract to limit liquid permeability through the PDMS device walls. Xenopus extract containing our soluble fluorophore was then pumped into the device at a rate of 7 μl/min for a total of 30 channel volumes in a 4°C cold room to prevent MT nucleation. A crossflow of Novak 7500 containing 2% surfactant (PicoSurf 1, Sphere Fluidics) was then pulsed into the device from the opposite outlet at a rate of 23 μl/min in 5-μl increments for a total of 15 μl (**Figure 1a, b**). The device was then transported on ice to a temperature-controlled room for imaging.

### Microscopy and imaging

All microscopy experiments were performed at 17°C, using a scientific CMOS (complementary metal oxide semiconductor) camera (Flash 4.0, Hamamatsu) mounted on an IX81 stand equipped with a spinning-disk confocal head (CSU-X1, Yokogawa). Confocal illumination was provided to the system by a LMM5 laser launch (Spectral Applied Research). Integration of all imaging system components was provided by Biovision Technologies. Image acquisition was performed using Metamorph 7.7 software (Molecular Devices). Images were acquired at the coverslip surface at 0.5-sec intervals for 1 min using Olympus objectives of varying magnification: 20× (0.85 NA) and 60× (1.35 NA) immersed in Olympus immersion oil (IMMOIL-F30CC). All time images were acquired within 20 min of the device being taken off ice. Multi-dimensional z-stack imaging of encapsulated extract allowed for precise determination of the encapsulated micro-enclosure diameter and height (as determined by the fluorescent signal), which were then used to calculate the volume of each micro-enclosure.

### Quantification of MT growth rates

MT growth rates and EB1 positional data were quantified using u-track particle tracking software (ver. 2.0; [24, 25]). Object tracking was done using “Microtubule plus-ends”, with “Comet Detection” being used to identify EB1 comets. The “High-pass Gaussian filter” and “Watershed minimum threshold” were determined in part by the NA of the objective, the bit depth of the camera, and the aspect ratio of the EB1 comets (typically, a High-pass Gaussian filter of 7 and a Watershed minimum threshold of 5 were used). For tracking, a minimum track length of three frames was kept constant across all treatments. The upper and lower bounds for the “frame-to-frame linking” were set at twice the mean growth rate in pixels per second of EB1 comets measured by manual kymographs for each specific treatment to reduce false-linking events. No forward reclassification was used in post-processing of tracks, and tracks starting in the first frame and those ending in the last frame were removed for volume and density comparisons (**Figures 2, 3**) Note that these tracks were included in the local-density analysis for ease of indexing and coding (**Figure 4**). All parameters set for automated detection and tracking were evaluated against a reference frame, in which automated detection of EB1 comets was compared to manual detection for a representative area. A 95% accuracy rate (for both false-positive and false-negative detections) was deemed sufficient for this analysis. MT growth rates were also measured manually for all treatments using kymographs (**Supplementary Figure 3**).

### Statistical analysis

Indicated statistical tests were conducted using Igor Pro 7 software (ver. 8.0.3.3; Wavemetrics) and confirmed in Excel. For each condition, measurements were taken from at least three independently prepared extracts.

## Acknowledgements

We thank Tim Mitchison and Keisuke Ishihara for helpful discussions. We would also like to thank Priscilla Phan for preparation of Xenopus extracts, and Alexandre Matov for his consultation and time in getting our lab set up to use u-track. This work was made possible by an Institutional Development Award (IDeA) from the National Institute of General Medical Sciences of the National Institutes of Health under Grant # 2P20GM103432. It was also supported by additional funding provided by the NIGMS under Grant #R01GM113028-and the Biomedical Scholars program of the Pew Charitable Trusts.

