## Supplemental Figures for "Microtubule growth rates are sensitive to global and local changes in microtubule plus-end density"

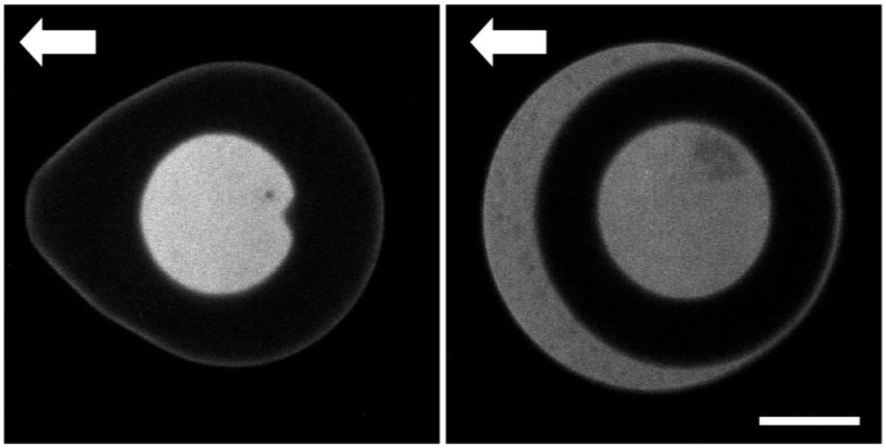

**Supplementary Figure 1. Isolating discrete volumes using oil crossflow.** PEGDA micro-enclosures under oil crossflow (direction of oil crossflow is indicated by the white arrow). Fluorescent signal (EB1-GFP) is trapped on the exterior of the micro-enclosure when a circular PEGDA structure (right image) is subjected to oil crossflow**,** whereas the fluorescent signal is contained within tear-drop shaped micro-enclosure (left image; scale bar = 50 μm).

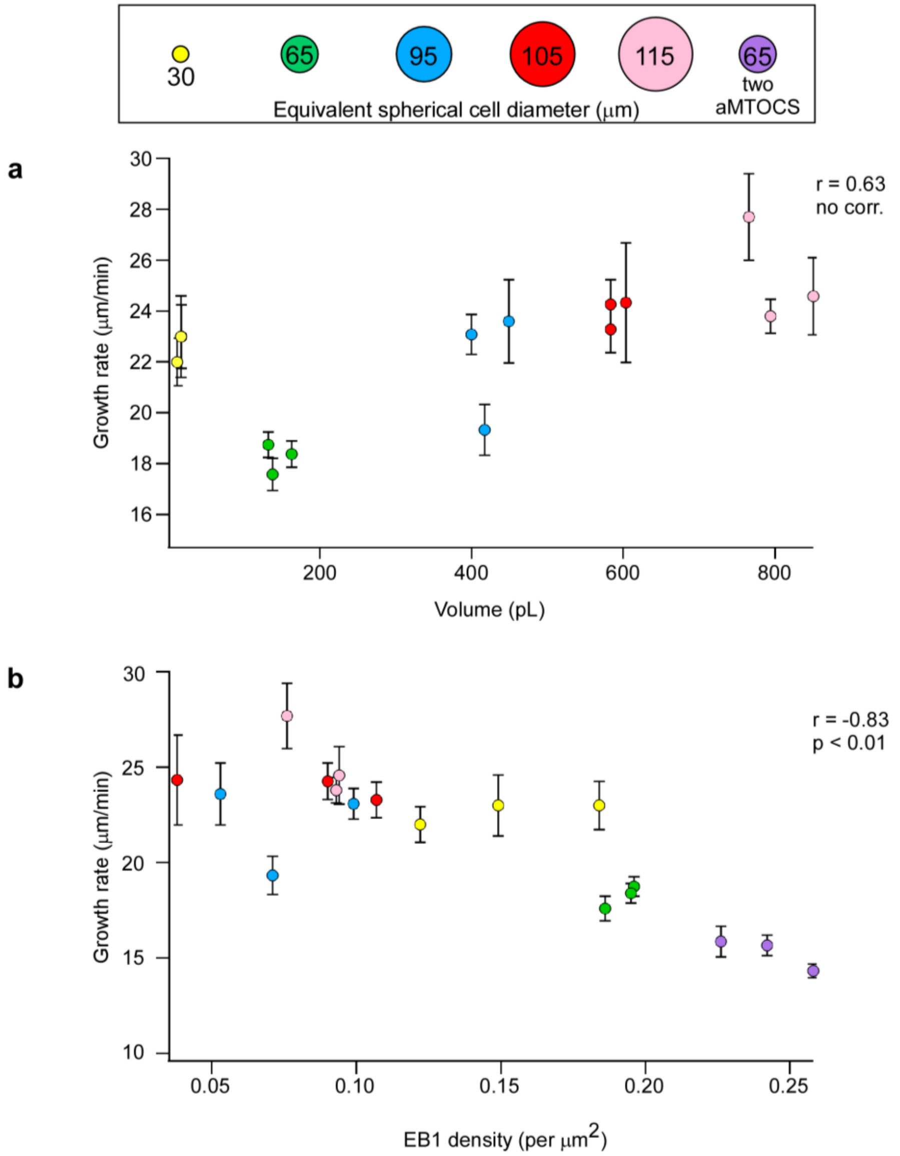


**Supplementary Figure 2. Manual measurement of MT growth rates. (a)** MT growth rates, as determined by manual kymographs, plotted as a function of cytoplasmic volume (n ≥ 30 kymographs per data point). Error bars equal two SEMs. Pearson correlation coefficient (r) is indicated at the side of each graph and is significant if $p < 0.01$ (the absence of a correlation is indicated otherwise). **(b)** MT growth rates from manual kymographs (a) plotted against EB1 density.

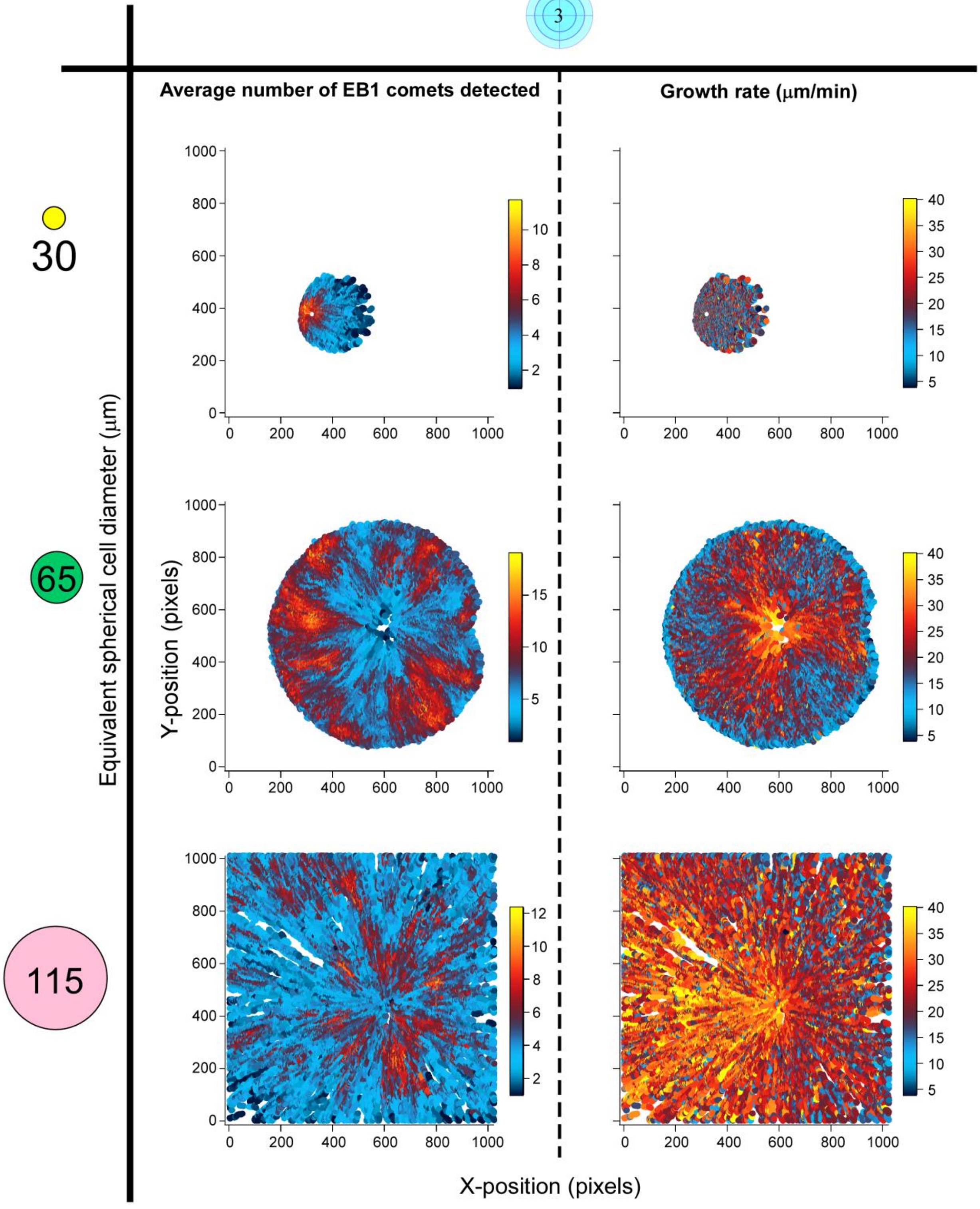


**Supplementary Figure 3. MT plus-end density and MT growth rates as a function of position.** The average number of EB1 comets detected within a 3 μm search radius and the average growth rate of each EB1 comet plotted against coordinate position for micro-enclosures of ~11, ~160, and ~800 pL (spherical cells of 30, 65, and 115 μm Ø, respectively).

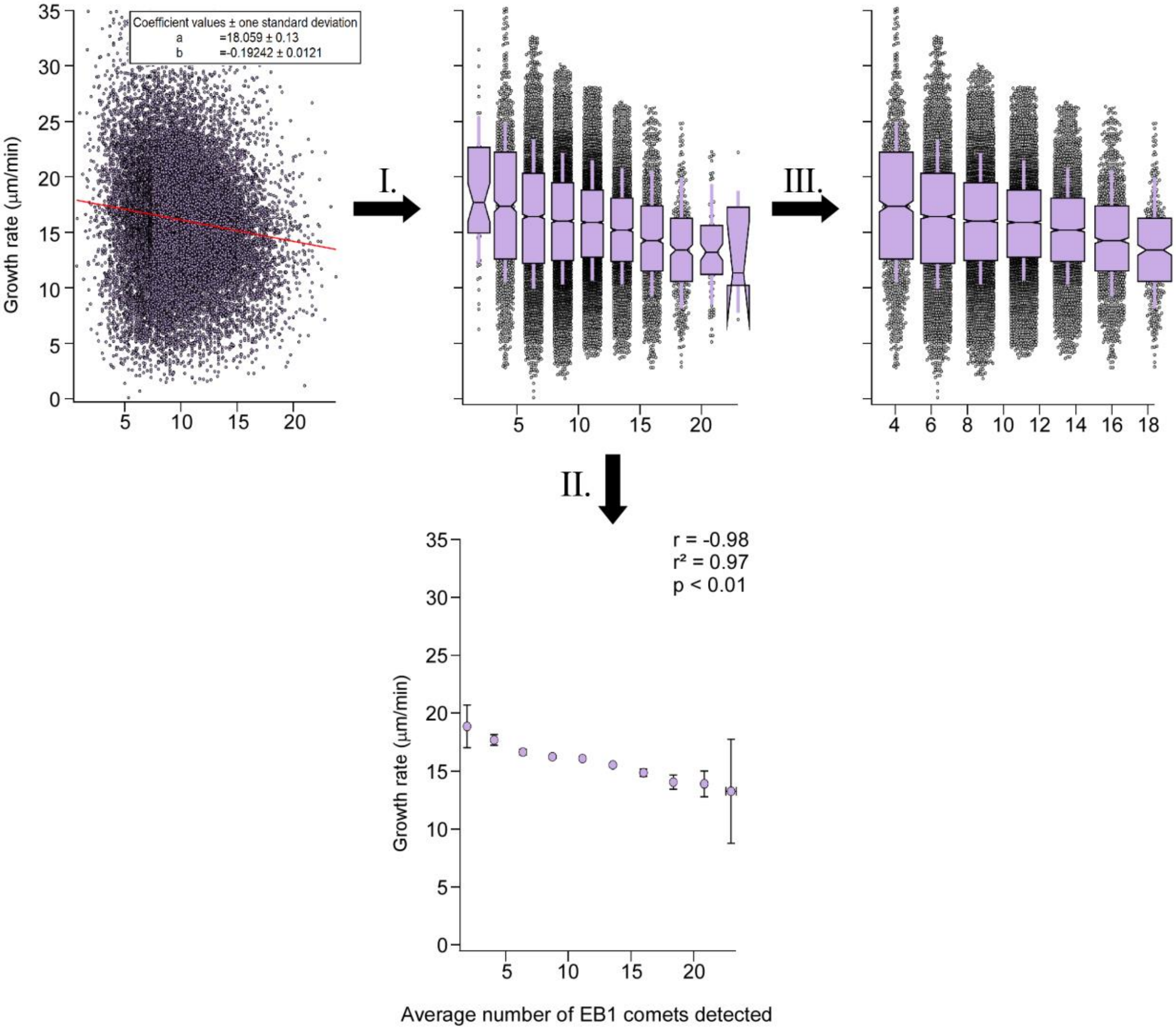


**Supplementary Figure 4. Local MT plus-end density analysis workflow.** Average velocity (i.e., growth rate) for every EB1 comet plotted against its local density (average number of EB1 comets detected in a 3 μm search radius). Every dot corresponds to a single EB1 track (n = 20,481). The slope of the spread was determined using a linear fit, with y = a + bx. **(I)** The raw spread from the analysis was subdivided into 10 bins, with each bin consisting of a ~10% increment of the highest local density observed for that search radius. In this example, the highest local density observed was 25 EB1 comets within a 3-μm search radius. As a result, each bin spans a range of approximately $0.1 \times 25$, or 2.5 EB1 comets, with each bin centered on the mean local density contained within the bin. **(II)** The mean local density and mean growth rate of each bin was then used to determine Pearson's correlation coefficient (r), Pearson's coefficient of determination ($r^2$), and the p-value for each local density analysis. All statistical analyses were performed on the entire data set (II). **(III)** Each bin was then plotted as a box plot, with Tukey-style IQRs, whiskers displaying one SD, and notches indicating the 95% confidence interval of the median. For graphical display (**Figure 4**), those bins containing <1% of the total population (for this example, 205 data points) were removed.

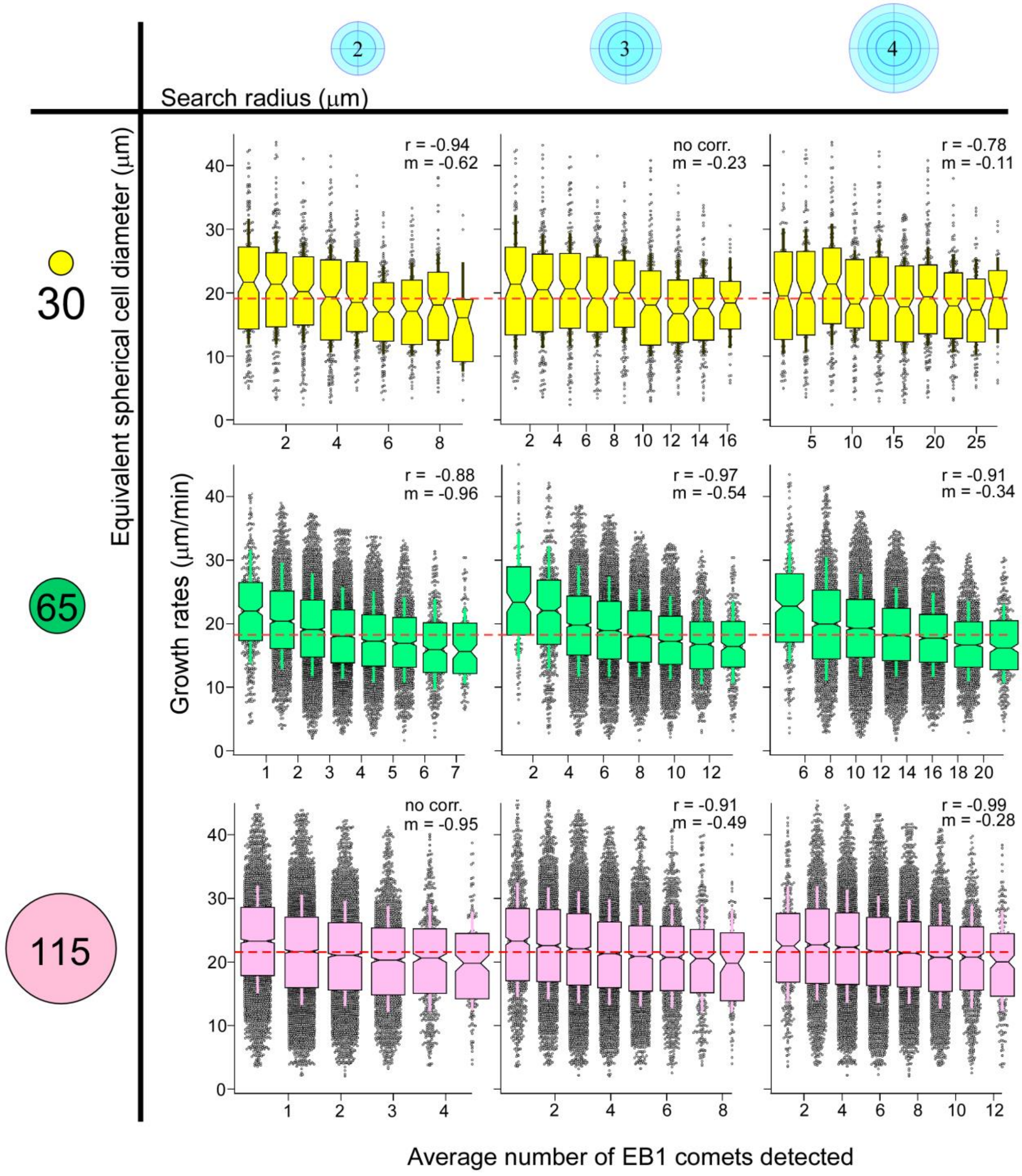


**Supplementary Figure 5. Local MT plus-end density across cytoplasmic volumes.** Average MT plus-end densities as a function of search radius (indicated by blue circles) for micro-enclosures of ~11, ~160, and ~800 pL (spherical cells of 30, 65, and 115 µm Ø, respectively). Dashed red line indicates the mean global growth rate for the system. Box plots are displayed with Tukey-style IQRs, whiskers displaying one SD, and notches indicating the 95% confidence interval of the median. Pearson's correlation coefficient (r) is displayed at the top of each graph with the slope (m) of the linear fit. Correlations were significant if $p < 0.01$ (the absence of a correlation is indicated otherwise).
